# T-Rex: Standardized Analysis of Germline Variants in Whole-Exome Sequencing Trios

**DOI:** 10.64898/2026.03.30.715083

**Authors:** Sara-Luisa Reh, Carolin Walter, Judith Lohse, Tabita Ghete, Markus Metzler, Anne Quante, Julia Hauer, Franziska Auer

## Abstract

Whole-exome sequencing (WES) enables the identification of rare germline variants contributing to pediatric diseases. Trio-based sequencing, comparing affected children with their parents, is particularly effective for rare disease genetics. However, WES data analysis requires bioinformatics expertise, varies across institutions, and is often incompatible with clinical workflows.

We developed T-Rex (**T**rio **R**are variant analysis of **EX**omes), a cross-platform desktop application that enables the standardized and local analysis of WES germline Trio data without the need for programming knowledge. T-Rex integrates state-of-the-art tools for alignment, dual-variant calling (GATK HaplotypeCaller + VarScan2), annotation (SNPEff/SNPSift), rare-variant filtering based on population frequencies (gnomAD), and family-based statistical testing, including the Transmission Disequilibrium Test with multiple-testing correction. Benchmarking of the dual-caller strategy on the Genome in a Bottle Ashkenazim Trio demonstrates high precision (99.2%) while maintaining robust sensitivity (91.1%). User testing (n=13) confirmed quick learning across clinicians and researchers. Application to a cohort of n=121 pediatric cancer Trio datasets, filtering for rare protein-coding variants (MAF≤0.1% in gnomAD v4.0), validated all assessable previously reported pathogenic variants.

Overall, T-Rex enables clinicians to robustly analyze WES Trio data in compliance with data protection regulations without requiring additional software licenses. As one of the first platforms for comprehensive WES Trio analysis that requires no programming expertise while providing clinical-grade, end-to-end workflows, T-Rex facilitates collaborative research between clinics and reduces reliance on external providers.

**Implementation and Availability:** The source code is available on GitHub (https://github.com/SaraLuisaReh/trex). The fully precompiled app is available on Zenodo (https://zenodo.org/records/19135262).

## 1. Introduction

Rare diseases collectively affect millions of individuals, yet for over 5,000 conditions, the genetic etiology remains unknown (Ferreira, 2019; Haendel et al., 2020). Progress in rare disease genomics is constrained by the difficulty of recruiting sufficiently large patient cohorts, which in turn limits the generation of statistically robust results and the application of artificial intelligence approaches. As individual centers typically encounter few affected individuals, meaningful studies rely on collaboration across institutions. Such collaboration is also essential for the delivery of high-quality genomic services and improved patient care (Julkowska et al., 2017; Raza & Hall, 2017). However, germline sequencing data constitute sensitive personal data, as they include heritable information with implications for both individual privacy and familial relationships. Thus, legal and ethical constraints on data protection often prevent the sharing of raw patient-level sequencing data, making decentralized, local analysis essential (Specht-Riemenschneider & Radbruch, 2021).

Since approximately 75% of rare diseases manifest in childhood, Trio studies, which sequence affected children along with both parents, have become a cornerstone for identifying pathogenic variants (JARC – Joint Action on Rare Cancers, 2020; Wright et al., 2023). By directly comparing transmitted and non-transmitted alleles within families, trio-based sequencing both reduces false positives and controls for population stratification, making it well-suited for small, heterogeneous cohorts assembled through multi-center collaborations (Aydin Son, 2013; Mahamad & Prashanth, 2020; Uffelmann et al., 2021). This is particularly relevant for childhood cancer, which arises largely in the absence of lifestyle-related exposures, and for which germline genetic predisposition has already been implicated in up to 15% of cases (Byrjalsen et al., 2020; Kentsis, 2020).

Despite the increasing availability of whole-exome sequencing (WES), clinical scientists often lack access to standardized, easy-to-use tools for Trio WES analysis (Koboldt, 2020; Kulkarni & Frommolt, 2017). Existing WES analysis pipelines commonly depend on command-line interfaces, operating-system-specific dependencies, or container platforms such as Docker or Nextflow, which require substantial programming knowledge and are not optimized for Trio studies, limiting adoption in clinical settings (Auwera & O’Connor, 2020; Danecek et al., 2021; Di Tommaso et al., 2017; Merkel, 2014; Ye et al., 2015).

To address these challenges, we developed T-Rex (**T**rio **R**are variant analysis of **EX**omes), a standalone, cross-platform desktop application providing a standardized WES pipeline for Trio studies, including rare-variant filtering and statistical testing. T-Rex enables clinicians to securely perform local WES Trio analyses without programming knowledge or sharing raw sequencing data. We demonstrate its utility using a Trio cohort of n=121 children with cancer and their parents.

## 2. Methods

### 2.1 Software Architecture

T-Rex is implemented in Python with a Tkinter/CustomTkinter graphical user interface (Lundh, 1999; Schimansky, 2024) and follows a Model–View–Controller architecture. The backend executes integrated Bash scripts for alignment, variant calling, and annotation, ensuring reproducible analyses across macOS, Linux, and Windows using only free and open-source software (FLOSS). T-Rex is distributed as a standalone desktop application, enabling straightforward installation without complex dependencies, administrative privileges, or command-line interaction. The platform was designed with a strong focus on usability, informed by regular feedback from clinicians and scientists. The interface is deliberately kept simple, with analysis options minimized, and interactive elements restricted during analysis to prevent accidental termination. Extensive in-platform instructions further support users without bioinformatics experience.

### 2.2 WES Processing Pipeline

The WES analysis pipeline processes short-read, paired-end Illumina data from Trio studies to identify germline variants in accordance with current software recommendations and best-practice guidelines, with optional support for single-end data. Users can initiate the pipeline at the alignment, variant calling, annotation, or filtering stage, enabling the analysis of FASTQ, BAM, VCF, CSV, and TSV input formats. A flowchart summarizing the pipeline implementation and the specific software tools used is shown in **Figure 1**.

**Figure 1.**
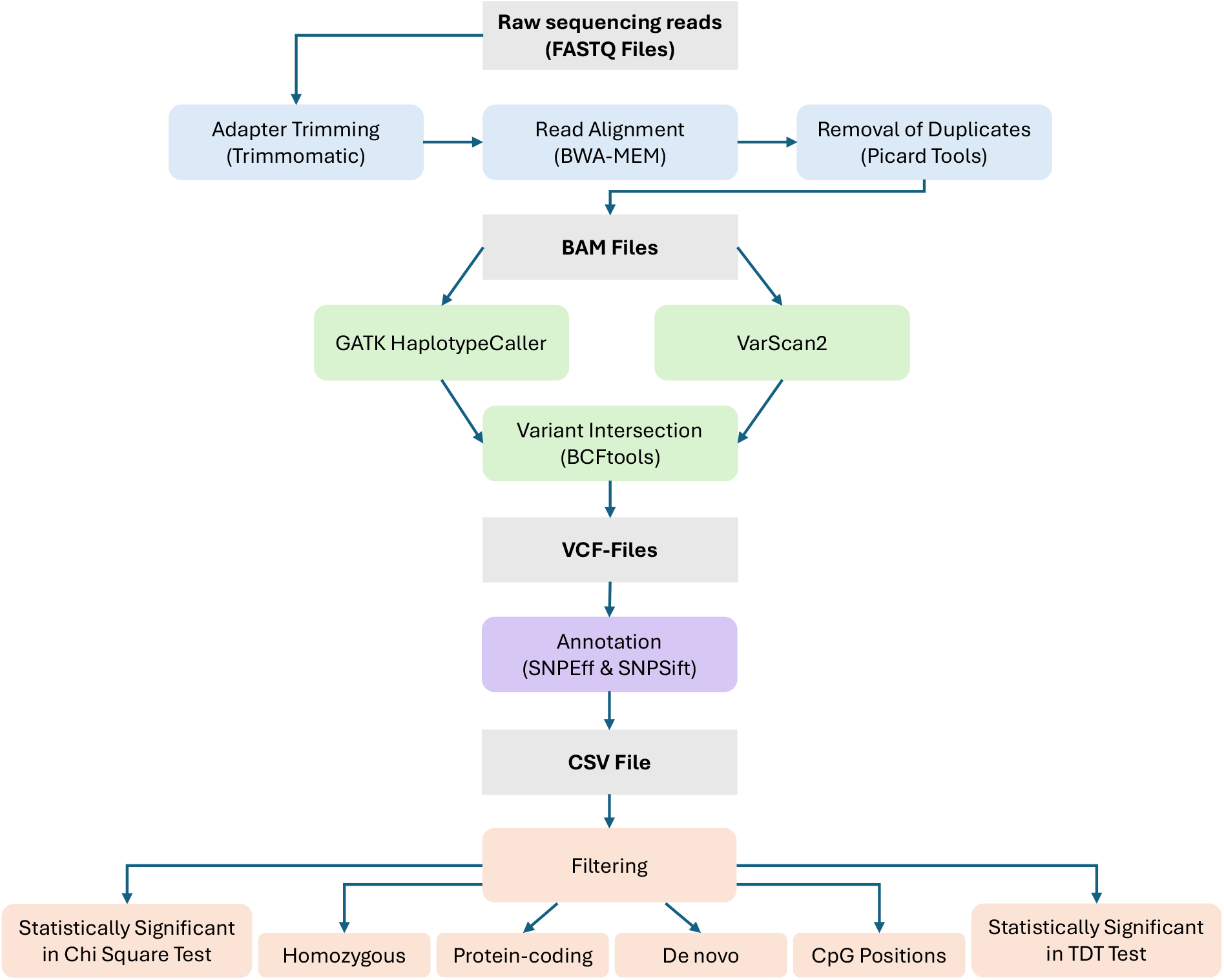
Flowchart illustrating the implementation of the WES data analysis pipeline.

#### 2.2.1 Pre-processing and Alignment

Following the current best practices for short-read alignment (Koboldt, 2020; Musich et al., 2021; Trevarton et al., 2023), Illumina paired-end reads undergo:

- adapter trimming with Trimmomatic (Bolger et al., 2014),
- alignment to GRCh38 primary assembly using BWA-MEM (Li, 2013),
- duplicate removal with Picard (Broad Institute, 2024),
- BAM indexing with SAMtools (Danecek et al., 2021).

#### 2.2.2 Dual Variant Calling

Following evidence that combining complementary variant callers improves accuracy (Bartha & Győrffy, 2019; Koboldt, 2020; Liu et al., 2017; Trevarton et al., 2023), T-Rex integrates two algorithms with distinct methodological strengths for germline variant detection. Since GATK HaplotypeCaller (Auwera & O’Connor, 2020) and VarScan2 (Koboldt et al., 2012) exhibit complementary performance characteristics (Liang et al., 2019; Liu et al., 2017), have both been validated for use with trio-based germline data (Koboldt, 2020; Lin et al., 2022), and have been recommended for ensemble use by prior studies (Koboldt, 2020; Liang et al., 2019), we implemented a dual-caller strategy integrating both tools for variant detection. Variant call sets were intersected using BCFtools (Danecek et al., 2021), retaining only variants independently identified by both callers to increase precision.

#### 2.2.3 Annotation and Filtering

Annotation is performed using SNPEff (Cingolani, Platts, et al., 2012) to predict functional impact, and SNPSift (Cingolani, Patel, et al., 2012) to retrieve allele frequencies from the Genome Aggregation Database (gnomAD) v4.0, including overall, European, and non-Finnish European populations (Broad Institute, 2023), as well as to assess variant pathogenicity based on ClinVar (Landrum et al., 2014).

After annotation, users can apply additional filtering options, including maximum allele frequency (default ≤1%) for a selected reference population, protein-coding variants only, homozygous or *de novo* variants only, variants located at CpG positions, and variants significant in either the Chi-Square Test or the Transmission Disequilibrium Test (TDT). When multiple filters are applied, only variants meeting all specified criteria will be included in the results.

#### 2.2.4 Statistical Testing

T-Rex implements two classes of statistical tests: (1) case–population comparison, using Fisher’s exact test when expected counts are ≤5 and Pearson’s χ^2^ test for larger counts (Fisher, 1934; Jaykaran, 2011; Pearson, 1900); and (2) case–parent comparison, using the TDT, which applies McNemar’s test or exact binomial testing based on transmitted versus non-transmitted alleles from heterozygous parents (He et al., 2014; Mahamad & Prashanth, 2020; McNEMAR, 1947; Spielman et al., 1993). Multiple-testing correction is performed using the Bonferroni method, as recommended for genome-wide rare variant analysis (Auer & Lettre, 2015; Ranganathan et al., 2016; Uffelmann et al., 2021).

### 2.3 Benchmarking with the GIAB Ashkenazim Trio

The dual-variant calling strategy was benchmarked on WES data from the Ashkenazim Trio reference samples (HG002 [child], HG003 [father], HG004 [mother]) provided by the Genome in a Bottle (GIAB) Consortium (Zook et al., 2016). Variant calling was performed using the integrated dual-caller workflow, combining GATK HaplotypeCaller v4 and VarScan2 in trio mode. Performance was assessed against the GIAB high-confidence truth set for HG002 (child), restricted to the exome target intervals. Precision, recall, and F1-score were calculated with RTG Tools v3.12 (Cleary et al., 2015). Benchmarking results from the consensus approach were compared to those obtained using GATK HaplotypeCaller v4 and VarScan2 individually, restricted to the same genomic regions.

### 2.4 Performance and User Testing

T-Rex was evaluated following standard software testing guidelines (Cohn, 2010; Pittet, 2024), including software, performance, and user acceptance testing.

Time and space complexity of individual pipeline components and the complete workflow were assessed on a system with 8 GB of RAM and 8 CPUs, using validated synthetic test sequences. In addition, wall-clock runtime and peak memory usage were measured on a system with 128 GB of RAM and 8 CPUs, using 121 real Trio WES samples.

User acceptance testing involved 13 participants (clinicians, research scientists, and doctoral candidates), aged 22–62 years (mean: 33.2 years), who completed structured tasks followed by semi-structured interviews, with task completion times recorded (**Supplemental Section 1**). Platform design was iteratively refined based on user feedback.

### 2.5 Trio Cohort Analysis

Analyzed germline DNA was derived from fibroblasts or blood and was previously sequenced at >100x coverage on Illumina NovaSeq (Friedrich et al., 2023). The raw FASTQ files were processed by applying T-Rex’s analysis pipeline, filtering for protein-coding variants with a minor allele frequency of less than 0.1% in the European reference population according to gnomAD v4.0 (n_reference_population_=582,716). Variant pathogenicity was assessed based on ClinVar annotations. The results were then compared with those reported by *Friedrich et al*. (2023) by evaluating whether the 14 variants classified as (likely) pathogenic by clinical geneticists in that study were also detected in the T-Rex analysis output.

### 2.6 Patient consent

The validation data is from patients <19 years of age, who were recruited unselectively at the Pediatric Oncology Department of the University Hospital and Faculty of Medicine Carl Gustav Carus, Dresden University of Technology (TUD) (years 2019 until 2021). Male and female participants were equally included without any selection bias for gender or sex. Consent of the families was obtained according to the Ethical Vote EK 181042019 (TUD).

## 3. Results

### 3.1 Linear Time and Constant Space Complexity

The time complexity across the pipeline was linear in cohort size (*O(n)*) (**Figure 2**), while the space complexity remained constant (*O(1)*).

**Figure 2.**
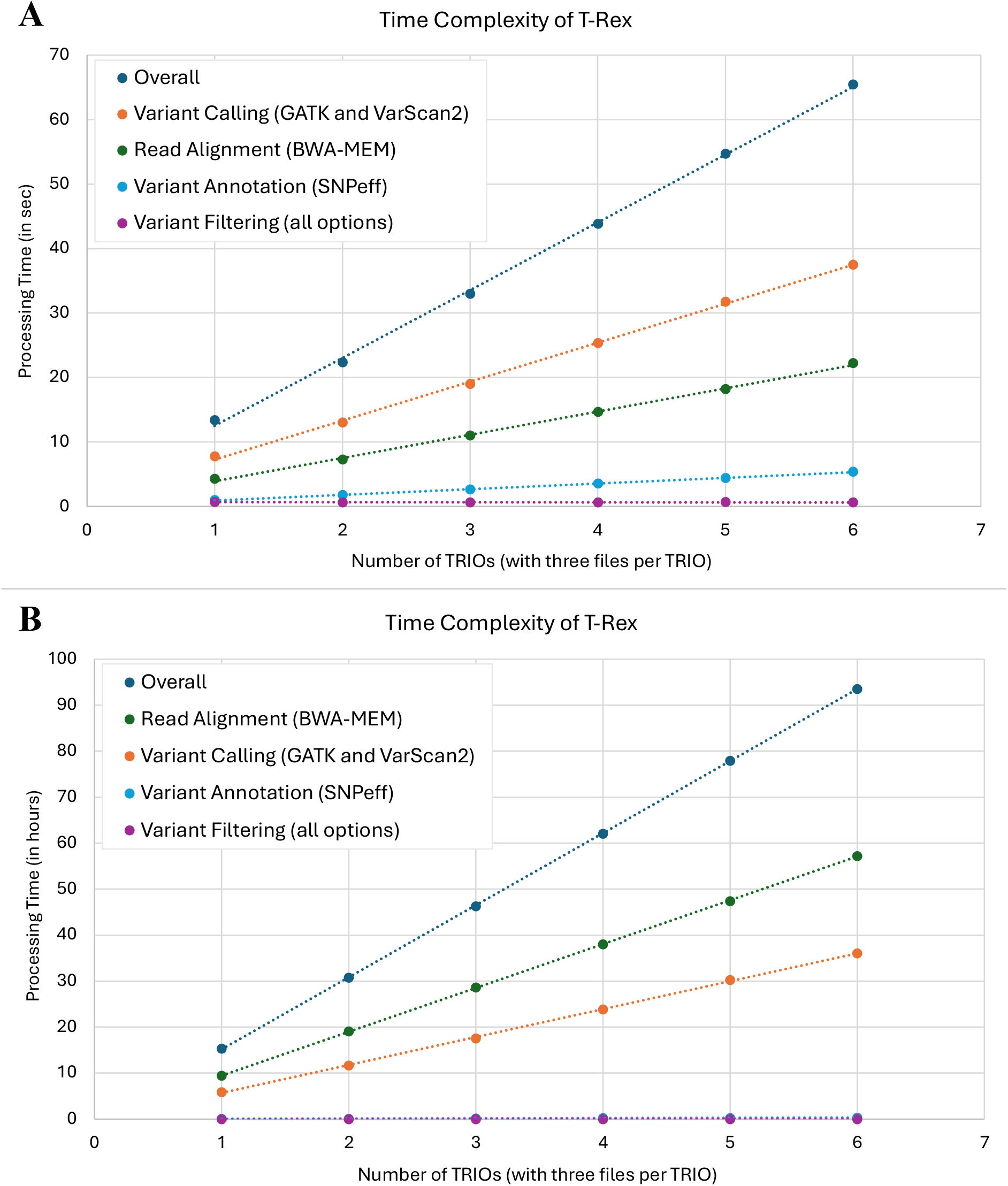
Time complexity of T-Rex on two different systems. A. Time complexity of validated synthetic test sequences on a system with 8 GB RAM and 8 CPUs. Variant calling with GATK HaplotypeCaller is slowed when RAM is below 16 GB, which explains why variant calling is slower than alignment on this system. B. Runtime of real WES Trio datasets measured on a server with 8 CPUs and a peak memory usage of 16 GB.

Memory usage adapted to system specifications but did not exceed 16 GB. When less than 16 GB of RAM was available, the runtime of GATK HaplotypeCaller increased, leading to longer overall analysis times, whereas the execution time of other pipeline components remained unaffected. On a server equipped with 8 CPUs and peak memory usage of 16 GB, processing 121 real-world Trio WES datasets required 15.3±0.7 hours on average.

### 3.2 Platform Operation is Easily Learned

All 13 participants learned to operate the platform in under 10 minutes, regardless of age or technical background. User feedback and iterative design adaptations further improved usability (**Supplemental Table S3**), enabling non-bioinformaticians to correctly and independently initiate Trio WES data analysis in under 2 minutes by the end of the study (**Figure 3**).

**Figure 3.**
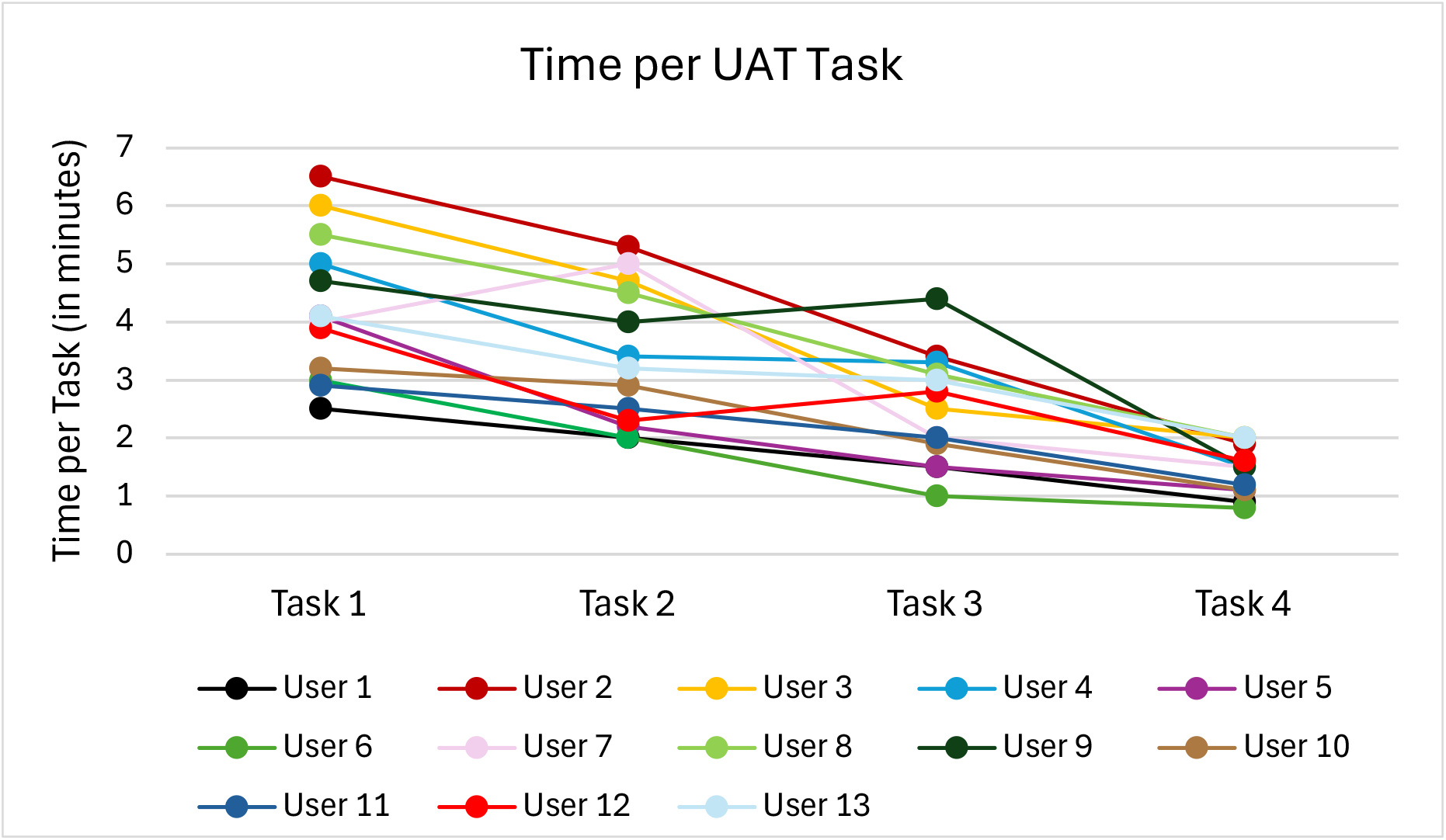
Comparison of task completion times during UAT across different users. Although task difficulty increased progressively, the time required per task decreased, indicating a learning effect.

### 3.3 High Precision of Dual-Caller Workflow on GIAB

To assess the accuracy of the dual-variant calling workflow, variant calls from WES data of the Ashkenazim Trio (HG002–HG004) were benchmarked against the GIAB high-confidence truth set for HG002 (child) within the WES target regions (Table 1).

**Table 1.**
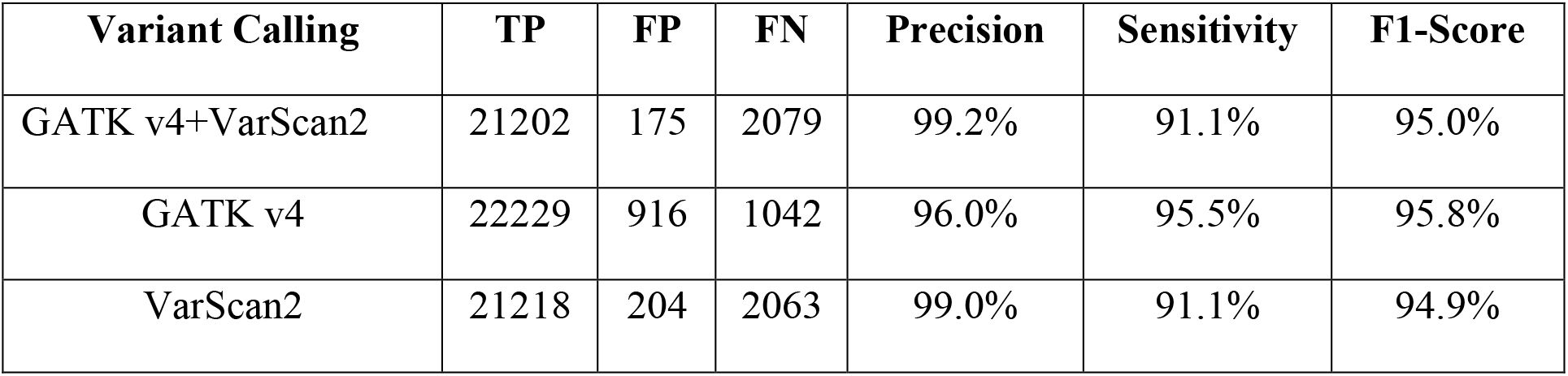
Performance of dual-variant calling on GIAB Ashkenazim Trio WES data for HG002 (child). Evaluation was performed in GIAB high-confidence regions intersected with WES target regions. True positives (TP), false positives (FP), false negatives (FN), precision, sensitivity, and F1-score are reported for individual callers (GATK HaplotypeCaller v4, VarScan2 in trio mode) and the dual-caller consensus workflow (GATK v4 + VarScan2). The dual-caller approach reduces false positives while maintaining comparable sensitivity. Note: In trio mode, only variants present in the child are reported, though parental allele information is included for downstream analysis.

The dual-caller consensus approach (GATK v4+VarScan2), which retained only variants identified by both tools, achieved a high precision of 99.2% with only 175 false positives while maintaining a sensitivity of 91.1%, resulting in an F1-score of 95.0%. Individually, GATK HaplotypeCaller v4 exhibited higher sensitivity (95.5%) but lower precision (96.0%) due to 916 false positives, whereas VarScan2 showed slightly lower precision (99.0%) but similar sensitivity (91.1%) with 204 false positives. These results demonstrate that the dual-caller strategy prioritizes high-confidence variant calls, substantially reducing false positives compared to the individual callers, while maintaining acceptable sensitivity, making it advantageous for clinical and research applications where precision in rare variant detection is critical.

### 3.4 Trio Cohort Characteristics

To validate the functionality of our platform, we re-analyzed published WES data from n=121 pediatric cancer Trios by *Friedrich et al., 2023*. The age at cancer diagnosis in the cohort (n=121) ranged from one month to 17.7 years, with a mean age of 7.1 years. The age at diagnosis is slightly bimodally distributed, with peaks in infancy (age 0 to 4) and adolescence (age 11 to 16) (**Figure 4A**).

**Figure 4.**
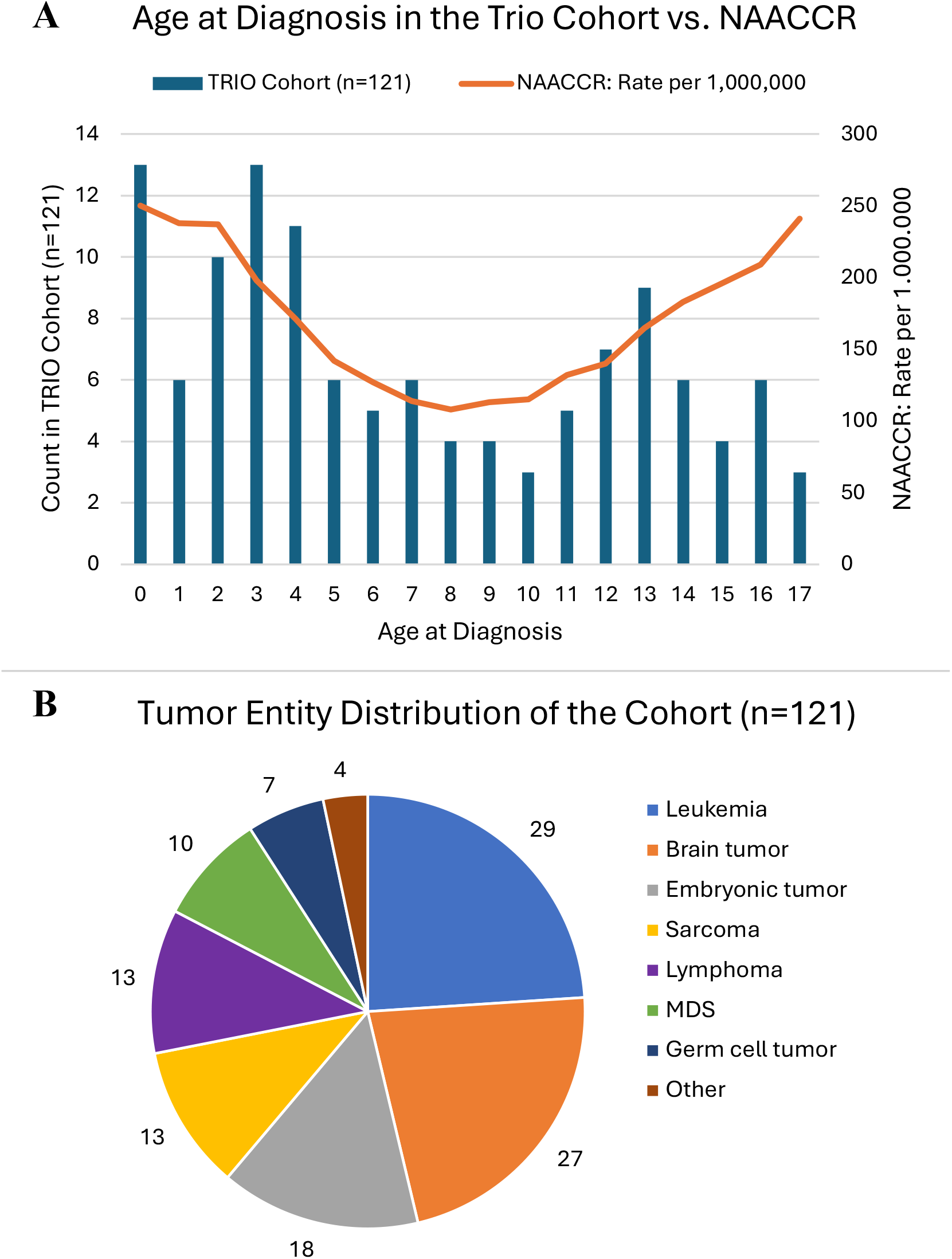
Characterisation of the Trio cohort. A. Comparison of the age at diagnosis of childhood cancer between the Trio cohort (n=121) and the North American Association of Central Cancer Registries (NAACCR) (based on Data from National Cancer Institute, 2024). The NAACCR documents the incidence of childhood cancer in 75% of all children in the USA (National Cancer Institute, 2024). B. Tumor entity distribution of the Trio cohort (n=121).

The most common cancer type in the cohort was leukemia (n=29), with the majority being B-cell acute lymphoblastic leukemia (n=22), followed by acute myeloid leukemia (n=4) and T-cell acute lymphoid leukemia (n=4) (**Figure 4B**). Brain tumors were the second most common cancer type in the cohort (n=27), followed by embryonic tumors (n=18), sarcomas (n=13), lymphomas (n=13), myelodysplastic syndrome (MDS) (n=10), and germ cell tumors (n=7). The “other” tumor category includes two melanomas, one carcinoid, and one case of histiocytosis.

### 3.5 T-Rex Detects Previously Reported Actionable Variants

First, we compared variant detection between T-Rex and *Friedrich et al. (2023)*. Of the 13 (likely) pathogenic variants assessable within our dataset, T-Rex detected all 13 (Table 2). Two additional variants reported by *Friedrich et al. (2023)* were not expected to be detected: one originated from a duo dataset (father-child) not included in our analysis, and one was an intronic variant located outside the WES target regions defined in the BED file.

**Table 2.** Comparison of variant detection between T-Rex and Friedrich et al. (2023). Of the 13 variants that were assessable within the dataset analyzed in this study, T-Rex successfully detected all 13. Two variants reported by Friedrich et al. (2023) were not expected to be detected: one originated from a DUO dataset not included in the analysis, and one was an intronic variant located outside the WES target regions defined in the BED file. These target regions can be expanded by the user if desired.

| Cancer Type | Gene | Variant | Effect | Detected by T-Rex |
| --- | --- | --- | --- | --- |
| MDS | ERCC6L2 | NM_001010895.2:c.1963C>T | stop_gained | ✓ |
|  | MECOM | NM_004991.4:c.2861G>C | missense | ✓ |
|  | WRAP53 | NM_001143990.1:c.1192C>T | missense | ✓ |
|  | WRAP53 | NM_001143990.1:c.904G>T | missense | ✓ |
|  | BRCA2 | NM_000059.3:c.6269del | frameshift | ✓ |
| Neuroblastoma | BARD1 | NM_000465.4:c.951_957del | frameshift | ✓ |
|  | BARD1 | NM_000465.4:c.1932_1933del | frameshift | ✓ |
| Brain tumor | BRCA2 | NM_000059.3:c.4214_4218del | frameshift | ✓ |
|  | ATM | NM_000051.3:c.3085dup | frameshift | ✓ |
|  | TSC1 | NM_000368.4:c.1963C>T | stop_gained | ✓ |
|  | NF1 | *Intronic variant → not covered by the WES target BED file* |  |  |
|  | TP53 | NM_000546.5:c.499C>T | stop_gained | ✓ |
| Leukemia | BRCA2 | NM_000059.3:c.3G>A | start_lost | ✓ |
|  | CHEK2 | NM_001005735.1:c.1229del | frameshift | ✓ |
|  | POT1 | *DUO only → not in analyzed Trio data set* |  |  |

### 3.6 Rare Variant Landscape in the Trio Cohort

Across the cohort of n=121 children with cancer, 33,020 protein-coding variants with a minor allele frequency of less than 0.1% in the gnomad European reference population were detected. Thereof, 13,510 variants were annotated in ClinVar: 1,399 were classified as benign, 3,606 as likely benign, 6,741 as variants of uncertain significance (VUS), 1,628 had conflicting interpretations, 66 were classified as likely pathogenic, and 70 as pathogenic (**Figure 5**).

**Figure 5.**
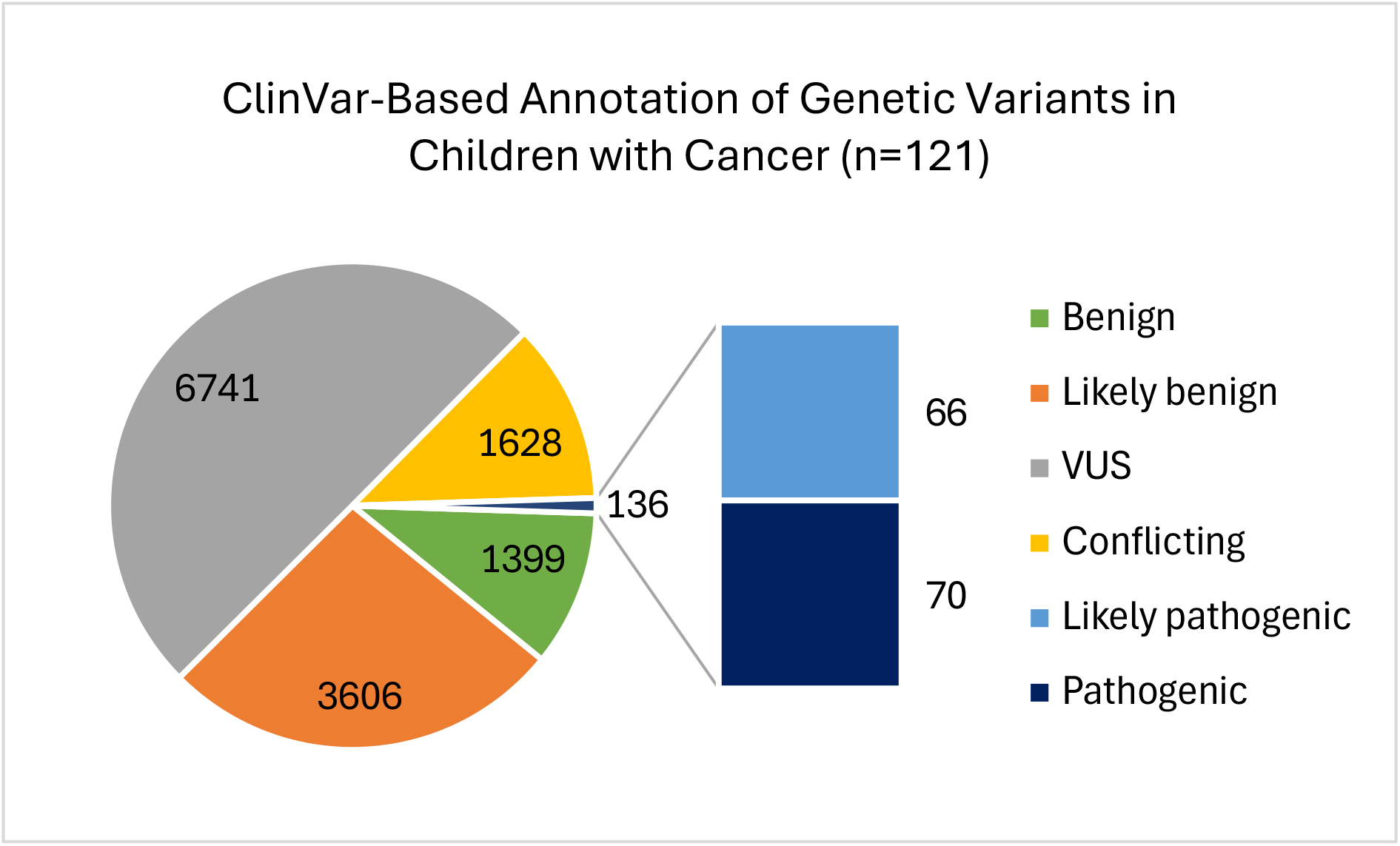
ClinVar annotation of rare variants (MAF≤1%) in n=121 children with cancer. VUS, variants of unknown significance. Conflicting, multiple different annotation labels in ClinVar.

Among the 70 pathogenic variants, 9 were annotated as cancer-associated in ClinVar. Of the 66 likely pathogenic variants, 3 were cancer-associated. The cancer-associated variants occurred in genes of the Fanconi anemia pathway (*FANCI, FANCD2, FANCM, FANCA*), in *SEC23B* (two variants, of which one is likely pathogenic), *ERCC6L2* (homozygous), *TSC1, WRAP53, BLM, MRE11* (likely pathogenic), and *ELAC2* (likely pathogenic) (**Table S4**).

Of these genes, only *TSC1* is associated with autosomal dominant inheritance and is therefore clinically relevant in the heterozygous state. In addition, the homozygous *ERCC6L2* variant was considered cancer-relevant. The child with the pathogenic autosomal recessive *WRAP53* variant carries an additional *WRAP53* variant on the second allele, currently classified as a VUS, which may likewise have actionable clinical implications. All remaining genes follow autosomal recessive inheritance and were not considered clinically relevant in heterozygous carriers.

These findings are fully concordant with the results reported by *Friedrich et al. (2023)*, as no additional pathogenic or likely pathogenic variants were identified by T-Rex beyond those previously reported. Accordingly, T-Rex achieved 100% sensitivity in detecting all assessable (likely) pathogenic variants from the reference dataset, while generating no false-positive pathogenic calls relative to the prior analysis.

## 4. Discussion

Family-based WES studies, such as Trio designs, are critical for identifying rare germline variants in pediatric diseases. However, current WES analysis tools are often fragmented into individual steps, require substantial programming expertise for operation, and often involve complex installation processes. (Auwera & O’Connor, 2020; Broad Institute, 2024; Danecek et al., 2021; Koboldt, 2013, 2020; Ye et al., 2015). Many of these packages must be operated via command-line scripts and rely on pre-installed software, posing challenges even for experienced bioinformaticians (Auwera & O’Connor, 2020; McLaren, 2024; Merkel, 2014). Containerized platforms such as Docker and Nextflow provide reproducible execution and prebuilt pipelines, respectively, but still demand programming skills and command-line operation, and Nextflow is limited to Unix-based systems (Di Tommaso et al., 2017; Docker, 2022; Merkel, 2014; Ribeiro-Dantas, 2025). JWES is cross-platform, yet analyses must be initiated via Java scripts, and the pipeline is restricted to single-sample data rather than Trio studies (Ahmed, 2021; Ahmed et al., 2021). In addition, although commercial tools for trio-based germline WES analysis are available, their high licensing costs, proprietary algorithms, and frequent reliance on external cloud infrastructures restrict their widespread adoption in academic and clinical settings.

T-Rex addresses these limitations as a unique Trio WES platform that operates entirely without programming knowledge, is fully cross-platform, can be installed and launched with minimal setup, and provides a standardized, reproducible, and user-friendly workflow.

This design enables decentralized analysis in settings where raw data sharing is restricted and bioinformatics support is limited, making WES analysis accessible to clinicians and researchers with diverse technical backgrounds while supporting the generation of large, harmonized cohorts across institutions — an essential prerequisite for the robust application of artificial intelligence approaches in rare disease research. Methodologically, T-Rex integrates established best-practice components for germline WES analysis with explicit support for family-based study designs. Unlike pipelines optimized for case-control cohorts, T-Rex incorporates trio-aware variant calling, inheritance-based filtering, and family-based statistical testing. Both case-population and case-parent statistical tests are included, allowing flexible analysis of rare variants across heterogeneous cohorts. Variant detection is performed using an ensemble strategy combining GATK HaplotypeCaller v4 and VarScan2, leveraging complementary strengths while intersecting calls to prioritize specificity and reproducibility, which is particularly important in rare disease studies. The performance of this dual-caller approach was benchmarked on the Ashkenazim Trio reference dataset from GIAB. Compared to the GIAB high-confidence truth set for HG002 (child), the dual-caller consensus workflow substantially reduced false positives (175 FPs) relative to the individual callers, while maintaining acceptable sensitivity (91.1%). In contrast, GATK HaplotypeCaller v4 alone achieved higher sensitivity (95.5%) but generated 916 false positives, whereas VarScan2 exhibited slightly lower precision (99.0%; 204 FPs) and similar sensitivity (91.1%). These results demonstrate that the dual-caller strategy effectively prioritizes high-confidence variant calls and reduces spurious calls, supporting its use in research and clinical contexts where precision in rare variant detection is critical. For studies where maximal sensitivity is required, the workflow can be configured to use a single caller (GATK HaplotypeCaller v4) rather than the dual-caller consensus.

Beyond its technical performance, T-Rex directly addresses the strategic objectives of national and international digital health initiatives, such as the German CORD-MI (Collaboration on Rare Diseases) project within the Medical Informatics Initiative (MII) and the European Solve-RD consortium (MII-Germany, 2024; Zurek et al., 2021). These programs aim to improve patient care by overcoming the fragmentation of genomic data across clinical centers. While these initiatives provide the legal and ethical frameworks for data sharing, T-Rex provides the necessary technical substrate for decentralized data harmonization. By enabling sites to execute identical, trio-aware pipelines locally, T-Rex facilitates a ‘federated analysis’ model consistent with GA4GH standards — where the analysis is brought to the data rather than the data to a central repository (Rehm et al., 2021). This approach ensures that even institutions with limited bioinformatics infrastructure can contribute high-quality, pre-processed variant calls to multicenter cohorts, fulfilling the mandate of the German Federal Ministry of Education and Research for privacy-compliant, cross-institutional research to improve diagnostic yields in rare disease populations.

In the here tested pediatric cancer validation cohort, the tumor distribution is closely aligned with the expected distribution based on the European Cancer Registry, except for nephroblastoma (n=3), which was slightly underrepresented in the study (n_expected_=6) (Kaatsch, 2010). Our analysis yielded ∼33,020 rare protein-coding variants, including 12 ClinVar-classified (likely) pathogenic cancer-associated variants. T-Rex successfully detected all previously reported (likely) pathogenic variants and did not identify additional pathogenic calls relative to the reference analysis, indicating 100% sensitivity and high specificity with respect to the published dataset, without compromising the detection of clinically relevant variants. User testing across different centers confirmed that T-Rex can be rapidly adopted by users with diverse technical backgrounds, and performance evaluation demonstrated linear time complexity and constant memory usage, enabling execution on standard local hardware.

T-Rex Limitations: T-Rex currently supports short-read Illumina WES and germline variant detection. Thus, whole-genome sequencing, long-read technologies, and somatic variant calling are not yet supported. The intersecting dual-variant-caller strategy prioritizes high-confidence calls but may reduce sensitivity, particularly for low-level mosaic variants or variants present only in one caller. For studies where maximal sensitivity is required, the workflow can be configured to use a single caller (GATK HaplotypeCaller) instead of the dual-caller consensus. Additionally, pathogenicity assessment relies on external annotation resources, which are subject to ongoing revision. Hence, final variant assessment must still be performed by a trained geneticist.

In conclusion, T-Rex is among the first locally executable, cross-platform WES analysis platforms specifically designed for Trio data that can be operated and installed without programming expertise. It integrates standardized, reproducible workflows with trio-aware variant calling and family-based statistical testing, accurately recovering known pathogenic variants without introducing additional false positives. The GIAB validation confirms the reliability of the dual-caller strategy for high-confidence variant detection, supporting its use in clinical and research applications. By removing technical barriers and enabling decentralized analysis, T-Rex facilitates collaborative studies across clinical settings and enables the discovery of rare pathogenic variants and the genetic mechanisms underlying disease in Trio WES data.

## Data availability

Variant pathogenicity annotations were obtained from the ClinVar VCF dataset (GRCh38, release 2024-01-03), available from the National Center for Biotechnology Information (NCBI). Allele frequency data were derived from gnomAD v4.0 exomes (GRCh38 assembly).

Whole-exome sequencing reference datasets for the Ashkenazim Trio (HG002, HG003, HG004), generated by the Genome in a Bottle (GIAB) Consortium, were downloaded from the European Nucleotide Archive (accessions SRR2962669, SRR2962692, SRR2962694).

The raw data sets generated and/or analyzed during this study are not publicly available because of personal data restrictions, but are available from the corresponding authors on request. This excludes any individual personal/ clinical data of the individuals, which would endanger their anonymity.

T-Rex is available as a pre-compiled application via Zenodo (https://zenodo.org/records/19135262), along with accompanying test datasets. The source code is publicly available at GitHub (https://github.com/SaraLuisaReh/trex). T-Rex is compatible with macOS (ARM64), Linux (x86; Ubuntu ≥24), and Windows (≥11 via WSL2).

