## Supplemental Tables and Materials for "T-Rex: Standardized Analysis of Germline Variants in Whole-Exome Sequencing Trios"

### Supplementary Figures and Materials

#### Supplementary Section 1

##### User Instructions for User Acceptance Testing

Please familiarize yourself with the platform. Read through the settings and the tasks below.

Setting:

You work in a clinic/research department that has sequenced the DNA of exomes from five children with cancer and their healthy parents. The raw sequencing data (FASTQ files) are stored on a hard disk. For a cooperation project with other clinics, you are asked to carry out the following tasks.

Tasks:

a) Use the FASTQ files and filter for all variants with a maximum allele frequency of 80% in the European reference population.

b) Use the CSV file produced in Task a) to filter for variants with a maximal allele frequency of 1%.

c) Use the BAM file to filter for only protein-coding variants with a maximal allele frequency of 5% in the Non-Finnish European population.

d) You have received Trio variant calling files (VCF Files) from a cooperating institute, and you are asked to filter for homozygous variants with a maximal allele frequency of 5% in the European population, which are significant in the Transmission Disequilibrium Test.

#### Supplementary Table S3

| Test person | Profession | Age | Task 1 | Task 2 | Task 3 | Task 4 |
|---|---|---|---|---|---|---|
| 1 | Research Scientist | 24 | ✓ | ✓ | ✓ | ✓ |
| 2 | Medical Doctor | 29 | ✓ | ✗ | ✓ | ✓ |
| 3 | Medical Doctor | 53 | ✓ | ✗ | ✓ | ✓ |
| Intervention: Replace the tab view, where the user can choose between multiple files and a single file, with a single input field, which automatically recognizes the type of entry. | | | | | | |
| 4 | Medical Doctor | 29 | ✓ | ✓ | ✗ | ✓ |
| 5 | Research Scientist | 31 | ✓ | ✓ | ✓ | ✓ |
| 6 | Medical Doctoral Candidate | 24 | ✓ | ✓ | ✓ | ✓ |
| 7 | Research Scientist | 24 | ✓ | ✗ | ✓ | ✓ |
| 8 | Medical Doctor | 58 | ✓ | ✓ | ✗ | ✓ |
| Intervention: Open the subsection, which lists the additional filtering options, from the beginning. | | | | | | |
| 9 | Research Scientists | 62 | ✓ | ✗ | ✓ | ✓ |
| 10 | Medical Doctoral Candidate | 22 | ✓ | ✗ | ✓ | ✓ |
| Intervention: Add a label on the top of the page that reminds the user to select the correct file format. | | | | | | |
| 11 | Research Scientist | 25 | ✓ | ✓ | ✓ | ✓ |
| 12 | Research Scientist | 24 | ✓ | ✓ | ✓ | ✓ |
| 13 | Medical Doctor | 27 | ✓ | ✓ | ✓ | ✓ |

*Table S3: User testing results and platform adjustments during user acceptance testing.*

#### Supplementary Table S4

| Cancer Type | Gene | Variant | Effect |
|---|---|---|---|
| MDS | ERCC6L2[a] | 9:95955996_C/T | stop_gained |
| | FANCI | 15:89315318_C/T | stop_gained |
| | FANCD2 | 3:10092206_G/A | stop_gained |
| | WRAP53[c] | 17:7701738_G/T | missense_variant |
| Leukemia | FANCM | 14:45166959_C/T | stop_gained |
| | ELAC2[b] | 17:13017778_G/A | missense_variant |
| Lymphoma | MRE11[b] | 11:94447276_G/A | stop_gained |
| Sarcoma | FANCA | 16:89799197_C/A | stop_gained |
| Germ Cell Tumor | SEC23B | 20:18515695_G/A | missense_variant |
| | SEC23B[b] | 20:18510875_C/T | missense_variant |
| Brain Tumor | TSC1[c] | 9:132905615_G/A | stop_gained |
| | BLM | 15:90761015_C/T | stop_gained |

*Table S4: Rare pathogenic variants identified in the Trio cohort (n = 121, MAF ≤ 0.1%) that are associated with cancer according to ClinVar. Allele frequencies were evaluated using the European reference population of gnomAD v4.0 (n = 582,716).*
*a Homozygous variant in the affected child.*
*b Variant annotated as likely pathogenic.*
*c Autosomal dominant.*
*In total, 12 (likely) pathogenic cancer-associated variants were identified. Thereof, three were cancer-relevant (denoted with a and c).*
*MDS, myelodysplastic syndrome.*
